# Crowding Effects in an Empirical Predator-Prey System

**DOI:** 10.1101/2020.08.23.263384

**Authors:** Sam Subbey, Anna S. Frank, Melanie Kobras

## Abstract

This paper uses a Lotka-Volterra (predator-prey) modeling framework to investigate the dynamical link between the biomass of an empirical predator, and that of its prey. We use a system of ordinary (ODE) differential equations to describe the system dynamics, and derive theoretical conditions for stability, in terms of system parameters. We derive the empirical system parameters by fitting the ODE system to empirical data, using non-constrained optimization.

We present results to show that the predator biomass is regulated by that of the prey. Furthermore, that the system dynamics is subject to Hopf bifurcation, conditioned on independent second-order terms in the ODE system. In ecological terms, the findings translate into evidence for existence of population crowding (density) effects.

## 1 Introduction

The classical Lotka-Volterra Model (LVM) [1, 2] defines a fundamental mathematical framework for studying predator-prey interactions. In ecology, innumerable studies on interacting populations have been based on varying formulations of the LVM. The LVM has therefore been referred to as the basic equation of ecology (see [3]). If *x* ∈ ℝ_≥0_ and *y* ∈ ℝ _≥0_ are state variables that represent population indices (e.g., abundance) respectively, of prey and predator, the LVM system, 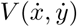, is defined by (1),

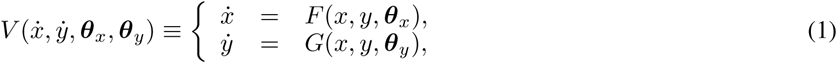

where *F* : ℝ_+_ *→*ℝ_+_ and *G* : ℝ_+_ *→*ℝ_+_ are continuous functions, and ***θ***_*x*_ ∈ ℝ^*n*^ and ***θ***_*y*_ ∈ℝ^*m*^ are sets of parameters associated with *x* and *y*, respectively. The literature contains several theoretical analyses [4, 5, 6] of how ***θ***_*x*_ and ***θ***_*y*_ determine the dynamic trajectory of the state variables.

If we follow, e.g., [7], in defining 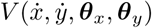 such that,

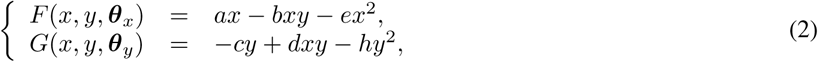

then ***θ***_*x*_ ≡{*a, b, e*}, and ***θ***_*y*_ ≡{*c, d, h*}. In (2), the term *ax* − *ex*^2^ defines a logistic growth with a limiting carrying capacity *a/e* (see e.g., [8]), and *bxy* is loss in prey biomass due to predation (also known as the functional response [9]. The predator dynamics is determined by a natural death rate term *cy*, population decrease due to intraspecific competition *hy*^2^, and *dxy*, which defines the biomass gain through predation.

Since the parameters in ***θ***_*x*_ and ***θ***_*y*_ determine trajectory of the state parameters, their definition is reflective of the underlying ecological assumptions. Thus, one may constrain any *θ* ∈ ***θ***_*x*_ (similarly for ***θ***_*y*_) to a scalar domain in R or to the domain of some continuous function in ℝ. For instance, the assumption that intraspecific competition is detrimental to population growth [10, 11] restricts both *e* and *h* to ℝ_+_. Seasonality is a characteristic feature of e.g., boreal and arctic environments [12, 13]. Because the authors in [14] assumed that this seasonality has continuous time effect, only the growth rates, both *a* and *c*, were defined as smooth sinus functions of time.

In practice however, constraining the domains of the model parameters may be excessively restrictive, especially when there is no empirical data to test validity of the ecological assumption. For instance, [15] presented a case where intra-specific competition, contrary to what is widely accepted, may have a positive effect on population dynamics. Hence, *e* and *h* are such that either *e* ∈ℝ _−_, *h* ∈ ℝ_−_ or *e*, ∈ *h* ℝ_−_. While seasonality may be an obvious driver of the system [16], incorporating such information in the population dynamics (of either predator and/or prey) may be non-trivial. This is because different functional expressions of the seasonality may lead to different scenarios of the population trajectory, see [17].

When data on both predator and prey are available, the parameters associated with *F* and *G* in (1) may be determined [18] by numerical optimization. If 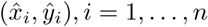, represent a set of empirical observations over *n* discrete time steps, and 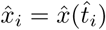 (similarly for *ŷ*), one may define the initial conditions for the ODE system by setting 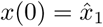, and *y*(0) = *y*_1_.

The estimation problem in (1) becomes even more challenging, when empirical data exists, only for either predator or prey, but not both. Then the derived trajectory of the missing component must be validated e.g., using auxiliary information to which it is linked (linearly or non-linearly).

This paper investigates the dynamics of a predator-prey system, where the empirical data available is that of the predator (not the prey), and measurements of the annual average weight of the predator. The predator is the Barents Sea (BS) capelin, which is a short-lived (1–4 years) fish species that spawn only once in their lifetime and die [19]. During the last three decades, the capelin biomass dynamics has been punctuated by several episodes of extreme biomass decline (collapse). These episodic events have been attributed to large predation (top-down effect) from other species, during crucial capelin life stages [20, 21]. The fact that food availability may also have regulatory (bottom-up) effect on the population dynamics has also been reported [22], but considered to be of less relevance than the top-down effect. The goal of this paper is to investigate whether the capelin biomass dynamics may be reconstructed (including episodic events of extreme decline) by solely considering a bottom-up regulation process.

The paper adopts the model definitions in (2), and uses unconstrained optimization to estimate the parameters of the system. We make inference about the system dynamics using the derived parameters.The manuscript is organized in the following way. Section 2 gives brief background information about capelin in the BS, and summary of the observation data on which the modeling is based. Section 3 revisits the mathematical models, and presents their theoretic dynamical system analyses. This section also presents a formal definition of the optimization problem, whose solution yields the system parameters. The theoretical results inform inference on the predator-prey behavior, on the basis of the derived parameters. Section 4 gives an overview of the numerical experiments, whose results are presented in Section 5, and discussed in Section 6. Our conclusions and discussion about possible limitations of the results, are presented in Section 7.

## 2 The Barents Sea capelin

Capelin is a focal forage species in the BS ecosystem. It preys on the lowest level species (zooplankton species), which are produced locally or are advected into the BS from adjacent areas [23, 24]. The selected prey size and type is age-dependent [22]. Capelin is itself, prey to other species (other fish and sea mammals) at higher levels in the food chain, and preferred prey for the Northeast Arctic cod [25].

Capelin has undergone several episodes of population collapse (defined as total biomass less than 10^6^ tonnes [19]) in the last three decades. The collapses have been attributed to inability of the stock to replenish itself (recruitment failure), resulting from high levels of predation (top-down effect) during crucial life stages [20, 21, 26, 27, 28, 29].

There is however, literature to suggest that food availability may have regulatory (bottom-up) effect on the population dynamics. Such effect has been reported for capelin in the Northwest Atlantic marine ecosystem off Newfoundland and Labrador, Canada [30]. The fact that the BS capelin may be subject to some degree of bottom-up regulation can be found in [22, 31], where the growth rate of young and older capelin were found to correlate, respectively, with the abundance of small and larger zooplankton. Other authors (e.g.,[24]) report on the link between changes in zooplankton biomass, and capelin biomass dynamics. Using statistical modeling and analysis, [32] showed that a bottom-up regulation capelin biomass dynamics is significant.

### 2.1 Data sources

Since 1972, the data on species abundance, spatial distribution and demography have been obtained from annual scientific cruises in the BS, during September [33]. The species abundance indices (length, weight, age, numbers) are converted into age-specific biomass.

Figure 1 shows the biomass of capelin from 1972 to 2013 (for the indicated ages), with notable stock collapses in 1985–1989, 1993–1997, and 2003–2006 [25]. This paper will use time series of the age-structured capelin biomass (Fig. 1) as predator data.

**Figure 1:**
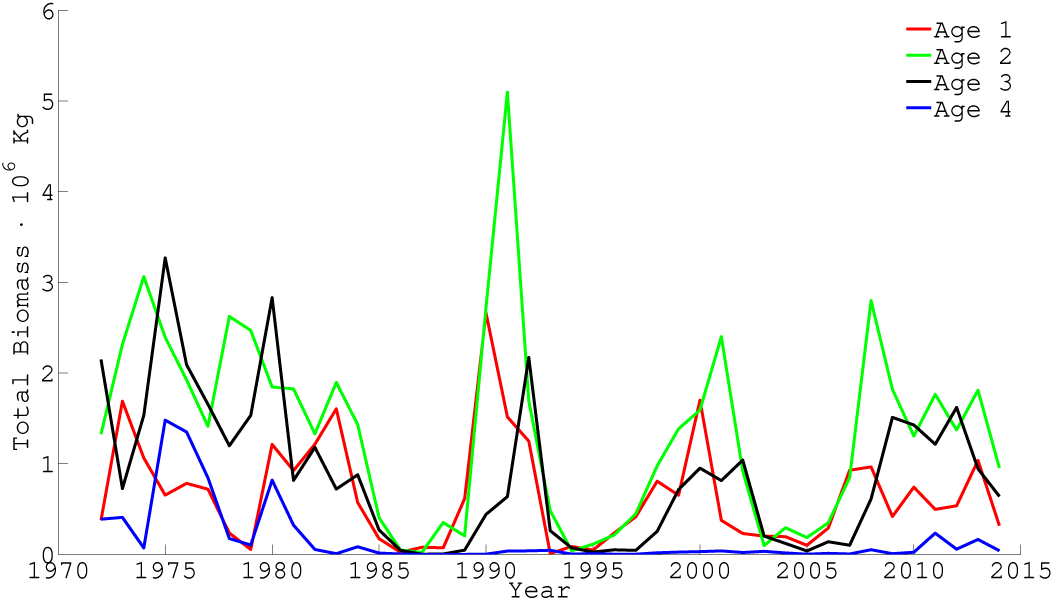
The capelin biomass data, obtained by an annual acoustic survey during the autumn

For parameter estimation and modeling of predator dynamics, we use a time-series of ages 1–3 capeling biomass data from 1990-2018. Data for age-4 capelin is excluded from the modeling and analysis because there are hardly any observations for this age-class in later years, and when observations exist, they are highly uncertain. Capelin biomass data are not reliable before 1983 [19], and capelin biomass was extremely low between 1986-1989. Hence our simulations (and parameter estimation) start in 1990, when the stock has recovered [34].

We have used the term *prey* to collectively describe all capelin prey, with a wide repertoire from Arctic copepod species to Atlantic krill [31]. This decision is influenced by the following considerations. Firstly, we have considered the prey data as unavailable partly because we are unable to pin down the exact prey type. Secondly, the intensive feeding season for capelin is July–October [33]. However, the scientific survey that collects data on the abundance of main capelin preys occurs at the end of this season. We consider this data therefore, as reflective of the residual prey abundance. The average weight of capelin has therefore been used as approximation for the validation of our derived prey biomass.

All data used in this manuscript have been obtained from the database of the ICES Working Group on the Integrated Assessments of the Barents Sea [35].

## 3 Mathematical model, Optimization and Analyses

This section presents the main mathematical model description, and the theoretical analyses.

### 3.1 The general predator-prey model

We use the model system defined in Section 1, where we model the dynamics of the capelin (*y*) and its prey (*x*) using the classical LVM system 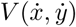. For *x* ∈ℝ_≥0_, *y* ∈ℝ_≥0_, *t* ∈ℝ_≥0_ we define

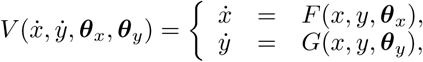

*F* : ℝ_+_ *→*ℝ_+_ and *G* : ℝ_+_ *→*ℝ_+_ are continuous functions, whose definition

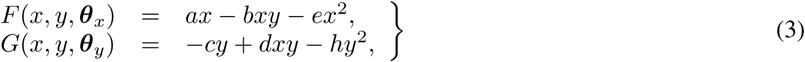

are adaptations from [7] and [36].

### 3.2 Theoretical analyses of system dynamics

#### 3.2.1 Coexistence and conditions

Our interest is in the non-trivial equilibrium point of the system 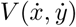, as the absence of either predator or prey is irrelevant for our quest. If we define *P*_∗_ ≡ (*x*_∗_, *y*_∗_) as the coexistence equilibrium point, we derive from (3) that

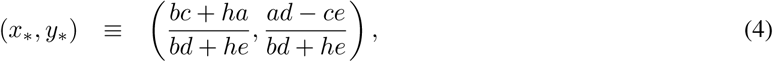

where

- Condition 1 (**C1**): *ad > ce*.

Shifting the equilibrium point (*x, y*) to the origin via the transformations *u*(*t*) = *x*(*t*) − *x*_∗_, *v*(*t*) = *y*(*t*) − *y*_∗_, results in (5).

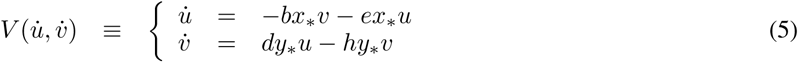

Let *p* = *ex*_∗_ + *hy*_∗_ ≥ 0, *q* = *hex*_∗_*y*_∗_ ≥ 0, and *r* = *bdx*_∗_*y*_∗_ ≥ 0, then (6) gives the characteristic equation for 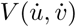, with discriminant *D* = *p*^2^ − 4(*q* + *r*).

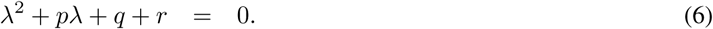

The equilibrium *P*_∗_ is stable if all roots of (6) have negative real parts Then the roots of (6) have (i) always negative real parts when {*p >* 0 ∧ *D* < 0}, (ii) always positive real parts when {*p* < 0 ∧ *D* < 0}, and (iii) at least one positive root for {*p* < 0 ∧ *D* ≥ 0}. We also define the following constraining conditions, which are applied later in this section:

- Condition 2 (**C2**): (*ex*_∗_ + *hy*_∗_) *>* 0 ∧ (*ex*_∗_ − *hy*_∗_)^2^ < 4*dbx*_∗_*y*_∗_,
- Condition 3 (**C3**): (*ex*_∗_ + *hy*_∗_) < 0.

#### 3.2.2 Stability and bifurcation analysis

Using the above conditions, we derive Lemma 1.

**Lemma 1**. *P*_∗_ *is stable for* **C1**∧**C2** *and unstable for* **C1**∧**C3**

□

Let tr𝕁(*P*_∗_) and D𝕁(*P*_∗_) define respectively, the trace and determinant of the Jacobian matrix 𝕁 (*P*), at *P* = *P*_∗_, i.e.,

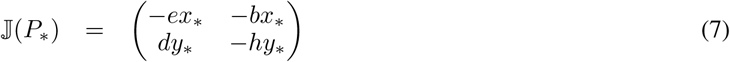

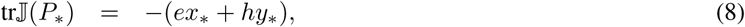

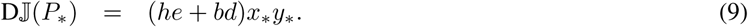

Observe that **C3** is a requirement that tr𝕁 (*P*_∗_) *>* 0, and that the trace depends only on the parameters *e* and *h*, which are respectively, coefficients of the second order terms *x*^2^, and *y*^2^. Since from **C1**, {*x*_∗_, *y*_∗_} ∈ℝ_+_, **C3** is only fulfilled when: (i) {*e, h*} < 0, (ii) *e* < 0 and |*ex*_∗_| *>* |*hy*_∗_|, or (iii) *h* < 0 and |*ex*_∗_| < |*hy*_∗_|.

Let **C1** prevail, we derive Lemma 2.

**Lemma 2**. *Let the 2-dimensional space* Ω *⊆*ℝ_+_ *×*ℝ_+_. *When* ***ζ*** = {*e, h*} ∈ Ω, *then* 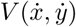 *undergoes no Hopf bifurcation at P*_∗_.

*Proof*. It is trivial to prove that

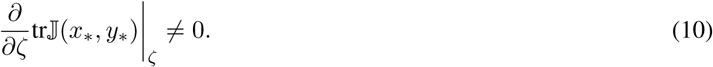

Assume ∃*h >* 0 for Hopf bifurcation at the equilibrium point *P*_∗_. When 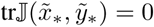, then *e* ≥ 0 iff *h* ≤ 0, which is a contradiction. The same argument applies when *e* is replaced with *h*. This completes the proof

□

Lemma 3 establishes Hopf bifurcation conditions for *e* and *h*.

**Lemma 3** ∃ *h*^*∗*^, *e*^*∗*^ ∈ℝ *and* ϵ ^*∗*^ *>* 0: 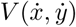 *undergoes Hopf bifurcation at P*_∗_ *for either of the following conditions*

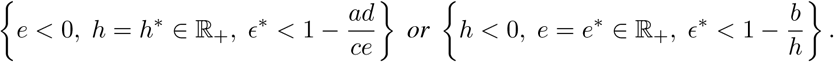

*Proof*. It follows from Lemma 2 that 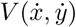 undergoes Hopf bifurcation under two conditions, namely, {*e >* 0∧*h* < 0} or {*e* < 0 ∧ *h >* 0}. We also prove that the condition 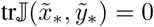 is satisfied by (11) for *h*, or by (12) for *e*.

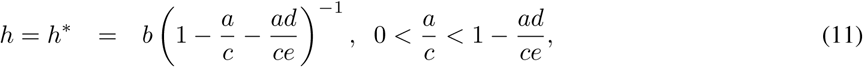

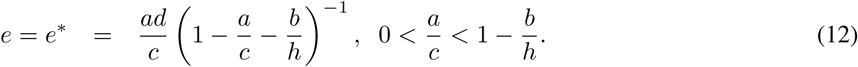

Define 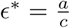, and note that *ϵ*^*∗*^ *>* 0, either when *e* < 0 in (11), or *h* < 0 in (12). This completes the proof

□

### 3.3 The optimization problem

Central to the optimization problem is the assumption that discrete empirical observations only exist for the predator, and not the prey. The goal of the optimization then, is to determine the set of system parameters that minimize the discrepancy between modeled and observed predator biomass.

Define {*i, n*} ∈ ℤ_+_ such that *i* = 1, …, *n*, and let 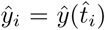 represent empirical observations of *y* over *n* discrete time steps 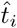. Note that *y* and *x* are coupled (through the functional response). Furthermore, since we assume no data exists on 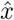, the initial condition *x*(0) must be estimated as part of the optimization procedure. The trajectory of *y* will also depend on *x*(0). Hence we write *y*(***θ***_*x*_, ***θ***_*y*_, *x*(0), *t*), and define Problem 1 as the general optimization problem.

**Problem 1**. *(The optimization problem) Determine* ***θ*** ≡ {***θ***_*x*_, ***θ***_*y*_, *x*(0)}:

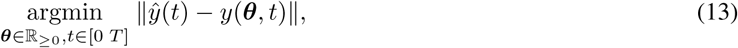

*where ŷ*(*t*) *is known at n discrete points at* 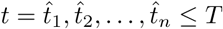.

## 4 Description of numerical simulations

Determining the trajectories of the predator-prey system combines the problem of integrating the system of coupled ODEs, and deriving the solution, ***θ***^*∗*^, of the optimization Problem 1. We use a numerical approach for integrating the ODE system, as for most of such coupled systems, finding closed form solutions is non-trivial [37].

For the coupled ODE system, we use the Matlab ODE solver, ode45, for numerical integration. The ode45 algorithm is based on RK5 (5th order Runge-Kutta) method for discretization, and RK4 for variable step selection. The initial values for the predator is taken to be the first empirical data point, i.e., *y*(0) = *ŷ*_1_, while *x*(0) = *x*_0_ is a parameter to be estimated.

The unconstrained model parameters are derived using the *fminsearch* algorithm in Matlab. The algorithm computes the unconstrained minimimum of given (linear/nonlinear) objective function using the Nelder-Mead algorithm (see [38]). For a predefined tolerance, the algorithm is considered to have converged after *J* iterations when the change in the objection function Δ*f* (***θ***_*j*_) <, for *j* ≤ *J*. For each candidate solution, ***θ***_*j*_ (*j* = 1, …, *J*), we obtain the spline-interpolated values of *y*(***θ***_*j*_, *t*) at *n* discrete (observation time) points 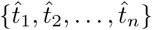. The objective function is then simply defined by (14).

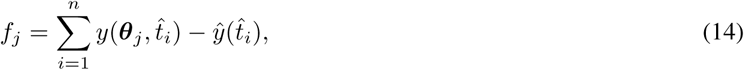

where, *f*_*j*_ = *f* (***θ***_*j*_). We obtain the system dynamics by analyzing the set of optimized parameters, based on the analysis presented in Section 3. We run the analysis for each age-class (1–3) of the predator.

## 5 Results from numerical simulations

This section presents results from our numerical simulations for the ODE model. We adopt the following notations, some of which has been used in Section 3, but repeated here for completeness sake:

- ***v*** = (*a, b, c, d, e, h*)
- ***v***^*o*^ ≡ optimized parameter set
- *e*^*∗*^ (similarly for *h*^*∗*^) ≡ critical Hopf bifurcation parameter
- *x*_0_ ≡ Initial condition for prey
- Let *e*^†^ (similarly for *h*^†^) define an arbitrary value such that:

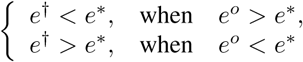

Table 1 summarizes the estimated model parameters and initial conditions for the predator-prey system for all age groups (1–3) of the predator. For age-2 capelin, the optimization algorithm fails to converge when the predator initial value *y*_0_ was assigned the age-2 capelin biomass at 1990, the highest ever recorded. Hence for age-2 capelin, our parameter estimation procedure used data from 1989-2018.

**Table 1:** The parameter estimation results.

| Age | $\mathbf{v}^o = (a, b, c, d, e, h)$ | $x_0$ | $e^*/h^*$ | $e^\dagger/h^\dagger$ |
| --- | --- | --- | --- | --- |
| 1 | (1.8308, 3.1757, 0.7462, 0.0730, 0.0150, -0.0245) | 7.5268 | $e^* = 0.0014$ | $e^\dagger = -0.0123$ |
| 2 | (0.1713, 0.1941, 2.3481, 0.4893, -0.0069, 0.2081) | 11.3527 | $h^* = 0.0316$ | $h^\dagger = -0.1449$ |
| 3 | (1.1394, 1.9504, 0.5666, 0.5627, -0.0097, -0.0497) | 2.4016 | $h^* = 0.0169$<br>$e^* = 0.0296$ | $h^\dagger = 0.0835$<br>$e^\dagger = 0.0689$ |

We use the optimized parameters to generate the predator-prey state variables. For brevity, we present results for age-3 capelin. Trajectories for other age-groups may be generated from Table 1.

Figure 2a.–b. (upper panel) show the simulated prey and predatory biomass dynamics, as well as comparison between modeled and empirical predator biomass data. The results show that the ODE system is capable of replicating the empirical observations. Figure 2c. shows the prey validation result. We see a consistent, temporal synchrony between the modeled prey trajectory and the averaged predator weight. Though the validation results for the age-1 and age-2 prey dynamics (see Fig. A.1) show identical trend with the scaled predator biomass data, the consistency in results (trend and level) appears to be degraded with decreasing predator age.

**Figure 2:**
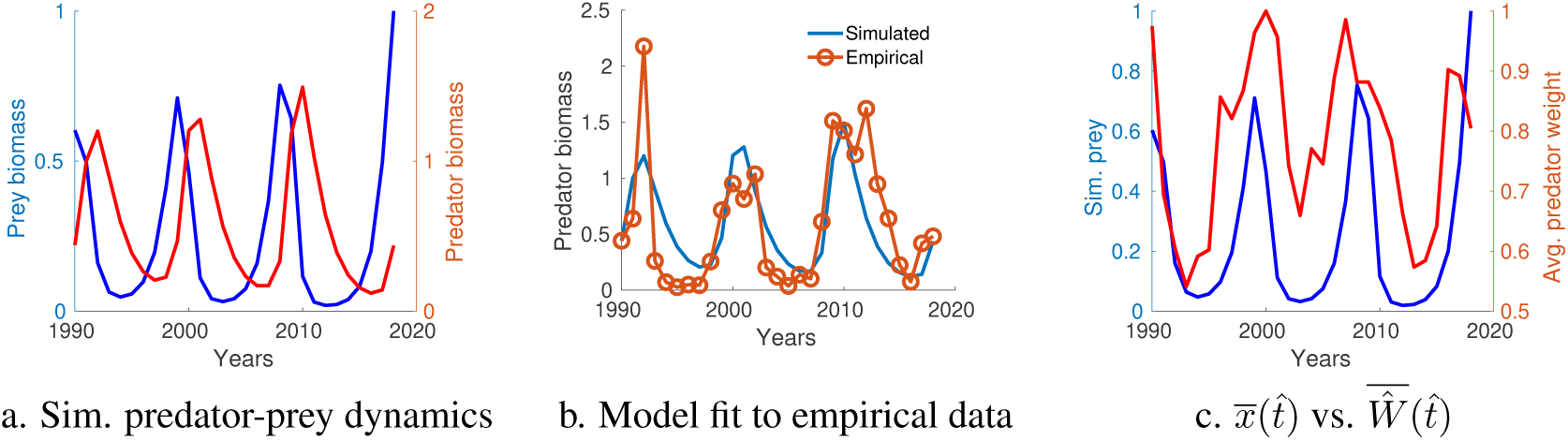
Optimization results for ODE problem – **Upper panel:** Simulated predator-prey dynamics (a.), model fit of predator dynamics to empirical data (b.) for age-3 capelin. **Lower panel:** Scaled total biomass of prey, 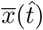, and scaled predator weight, 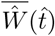, in the period 1990–2018 (c.)

Consistent with Lemma 3, results in Table 1 show that Hopf bifurcation point exists for parameters *e* or *h*. We examined the phase-plane dynamics of the system for both bifurcation parameters. Here, for brevity, we show the results for three scenarios of *h* at *h*^*o*^, *h*^*∗*^, and *h*^†^. The left panel of Fig. 3 shows model fit to data for the three scenarios, while the right panel shows the corresponding phase-plane dynamics of the predator-prey system. The phase-plane reveals a dynamic that is (a) Asymptotically unstable (*h*^*o*^) *→* (b) A limit cycle (*h*^*∗*^) *→* (c) Asymptotically stable (*h*^†^).

**Figure 3:**
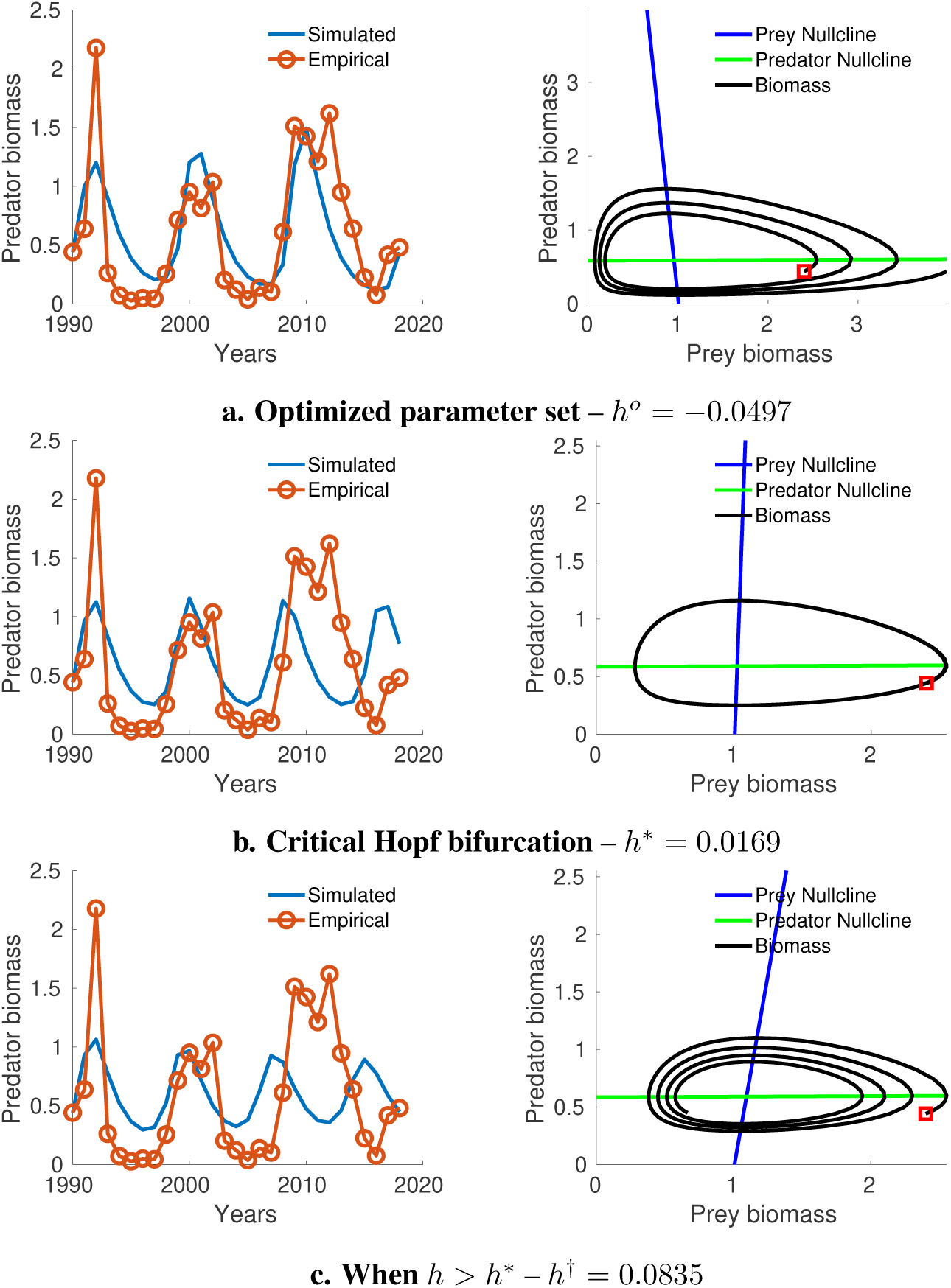
*h*-bifurcation analysis for ODE model – **Left column:** Model fit to data. **Right column:** Phase-plane plot of the predator-prey dynamics. Observe the phase-plane dynamic transitions from (a) Asymptotically unstable (*h*^*o*^) *→* (b) A limit cycle (*h*^*∗*^) *→* (c) Asymptotically stable (*h*^†^). The red square in the phase-plane plot is the origin of the time-series.

## 6 Discussion

We have used a Lotka-Volterra model formulations, to study the dynamics of an empirical predator-prey system, where data on the prey, prior to predator exposure, is absent. We have used auxiliary information about the predator to validate the simulated prey biomass dynamics.While full simulation results exist also for age-1 and age-2 capelin (Table 1), graphical results have only been presented for age-3. While this is partly for the sake of brevity, a stronger reason is that only the age-3 average capelin biomass weight appeared to be very consistent (in trend and level) with our modeled prey dynamics. For ages-1 and 2, though the trend in the modeled prey appears to be synchronous with the average weight of fish, the consistency is less pronounced, compared to that for age-3 fish. One plausible explanation of this observation could be that age-groups 1 and 2 are less affected by bottom-up effects compared to those of age-3. An alternative (and maybe stronger) argument may lie in the weight-at-age dynamics for the different age-classes.

Capelin feeding is size- (and therefore also age-) dependent. Small capelin eat small plankton organisms (small species or early life stages of larger species) while larger capelin are able to consume a wide range of crustacean plankton (copepods, euphausiids, amphipods, etc.) but prefer larger organisms (mainly euphausiids). The shift in diet is a gradual process but a major shift from copepods to euphausiids mainly happen in the second year of capelin life. In principle, the individual weight of age 1, 2 and 3 capelin may reflect the dynamics of various types of plankton, which may vary independently. The weight of age 1 capelin might be a good proxy for copepod abundance but not for euphausiid or amphipod abundance while weight at age 3 might be a proxy for euphausiid abundance or for an ensemble copepod/euphausiid/amphipod abundance [22]. Given the different age-dependent feeding scenarios, we investigated the ease with each total prey may be linked to the average weight for the different age classes. We investigated the variability in average weight when cohorts transition from one age-group to another.

If we define

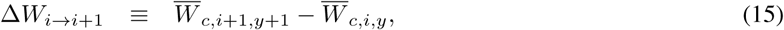

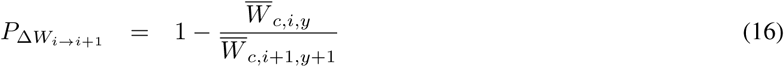

where 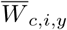 is the average weight of fish cohort *c* at age *i* in year *y* (and similarly for 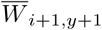), then for *i* = 1, 2, 3, we derive three groups of Δ*W*_*i→i*+1_. Figure 4 shows a Boxplot of data for the three groups of Δ*W*_*i→i*+1_, for cohorts from 1990-2008. This figure shows the highest variability in average weight from age 1 *→* 2 (approximately 68%), and lowest for 3 *→* 4 (approximately 15%). Thus the average weight at age-3 represents a near stable (asymptotic) weight of capelin. A stronger link may therefore, exist between the average weight and prey abundance. Consistent with Fig. 4, we expect this link (between prey abundance and average weight) to be poorer with decreasing age-group. These observations are consistent with our simulated prey abundance and average weight of fish (at age). Therefore for ages 1 and 2 capelin, a stochastic prey model (e.g., a stochastic differential equation) may be more appropriate for modeling the dynamics between predator and prey. The inherent stochasticity associated with ages 1 and 2 also implies that our simulated prey dynamics may be one of several stochastic realizations of the prey dynamics.

**Figure 4:**
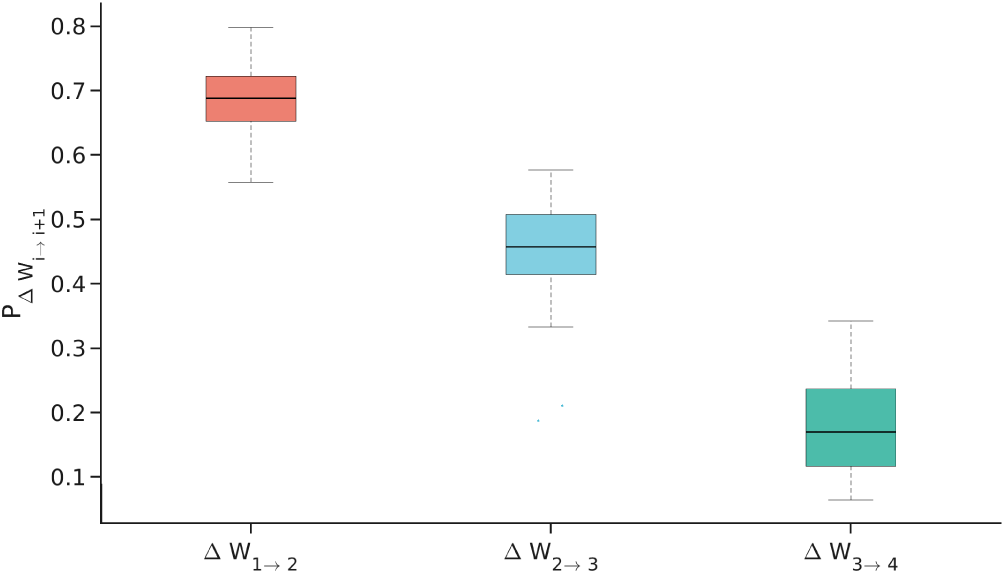
Boxplot of fractional change in average weight, 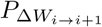, of cohorts at age *i* (in year *y*) and age *i* + 1 (in year *y* + 1) for *i* = 1, 2, 3., using data from 1990–2018.

There is a caveat in using average weight by age for fish of age 2 and higher, as the component of the stock that is measured is affected by the combination of length-dependent maturity and total spawning mortality. The largest age-2 fish measured in autumn, will mature, spawn, and die, before next autumn cruise. Consequently, these do not form a part of the cohort of age-3 fish whose mean weight are measured the following autumn. This may result in underestimation of annual mean weight for fish older than two years which, consequently, weakens the dynamic link between average weight and prey abundance. The underestimation (and its consequence) will be expected to increase with age (see [39]). On the other hand, since weight by age represents accumulated growth throughout the year up to the time of sampling, the accumulation process can potentially lead to a smoothness of the data uncertainty. Consequently, the link between food and weight at age becomes stronger (as our results indicate) with increasing age.

In general, our model shows that the biomass trajectories bifurcate at the critical point for negative coefficients ({*e, h*} < 0) of the second-order terms, i.e., *x*^2^ for prey, and *y*^2^ for predator. These parameters determine the stability state of the system dynamics (see Fig. 3). Our results (Table 1) show that for the age-classes, either *h* < 0 (age-1), *e* < 0 (age-2) or {*e, h*} < 0 (age-3). Higher population densities may lead to food competition that affect fish growth, and subsequently, delay reproduction and population growth [10, 11]. However, if we restrict the interpretation of the second order terms (involving *e* and *h*) to represent *intra-specific* (within species) competition, then according to the literature (see e.g., [40]) {*e, h*} ≥ 0. That a negative relationship exists between population growth and population density (also known as direct density dependence), is an *a priori* assumption in most models of harvested populations [15]. Hence, with a few exceptions (e.g., [15]), there is hardly literature on positive effects of high population densities on fish biomass growth.

The results show that labeling the second order terms as intra-specific competition may be restrictive. Define the second order terms by continuous functions, *C*(*x*) ∈ *C*^*α*^ and *D*(*y*) ∈ *C*^*α*^ (*α* ≤ 2), respectively for prey and predator. Then the ODE system can be written as

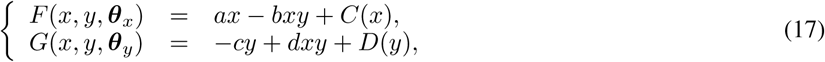

and the functions (*C*(*x*) and *D*(*y*)) may be interpreted more generally, as representing the regulatory effect of population density (crowding) on growth. The authors in [41] doubt that intraspecific competition may be a main determinant of capelin ecological condition. However, density effects include other processes in addition to growth, such as, survival, movement or dispersion (see [42]). Further to this consideration, is the fact that, due to their confounding nature, it is non-trivial to define the specific density effect(s) that regulate the dynamics of a fish population [43].

Our findings suggest that the signs of *h* and *e* determine the type of population regulation, which also has an effect on the stability of the predator-prey system. From Fig. 3, the predator-prey system is unstable for *h* < *h*^*∗*^ (similarly for *e*), but becomes stable when *h > h*^*∗*^. In other words, mortality resulting from high population density stabilizes the predator-prey system. Density effects that contribute to an increase in growth rates are destabilizing to the system. This demonstrates the regulating effect of density effects in capelin, even in the absence of fishing. The ability of our model to generate different dynamics, e.g., from damped oscillations to stable limit cycles, is a characteristic of mathematical models of population dynamics that include a density-dependent term (see [44]).

It is interesting to note that for age-3 capelin, the combination of prey and predator density effects (*h* and *e*) determines the dynamic stability of the system. The bifurcation in the dynamics of age-1 capelin results from prey density (*e*), while for age-2 capelin, it is solely the predator density (*h*) that dictates Hopf bifurcation.

## 7 Concluding remarks

Our modeling results give insight into the regulatory effects of prey on the BS capelin biomass dynamics. The results show consistency between our theoretical analyses, and simulations. We have furthermore validated the simulated prey dynamics by showing it to be perfectly synchronous (albeit with a slight time delay) with the annual average predator weight. Our results show that the stability and dynamic trajectories (of both predator and prey) are highly regulated by effects of population crowding (also referred to as density effects), though we are unable to pin down the specifics of these effects.

A key ecological highlight of our findings is that the biomass dynamics for ages 3 and older capelin may be regulated by a bottom-up effect, while this regulation is more difficult to quantify for younger ages (1 and 2). This is either due to the stochasticity in prey dynamics for younger fish, or that top-down (mortality due to predation from other species) effects dominate.

Finally, we have desisted in extending our inference to whether the identified regulatory effects (in isolation) explain the episodic collapses of capelin biomass. Such inference, in our opinion, must be based on integration of information in this paper, and analysis of other biophysical information in space-time. Unfortunately, such an undertaking is beyond the scope of this paper, and will therefore, be pursued in a sequel manuscript.

## Acknowledgements

SS is grateful to the Fulbright Foundation (Norway) for funding his research at Cornell University, where work on this manuscript was initiated. The authors are especially grateful to H. Gjøsæter (Institute of Marine Research, Bergen, Norway), for proofreading the ecological aspects of this manuscript.

## A Appendix

**Figure A.1:**
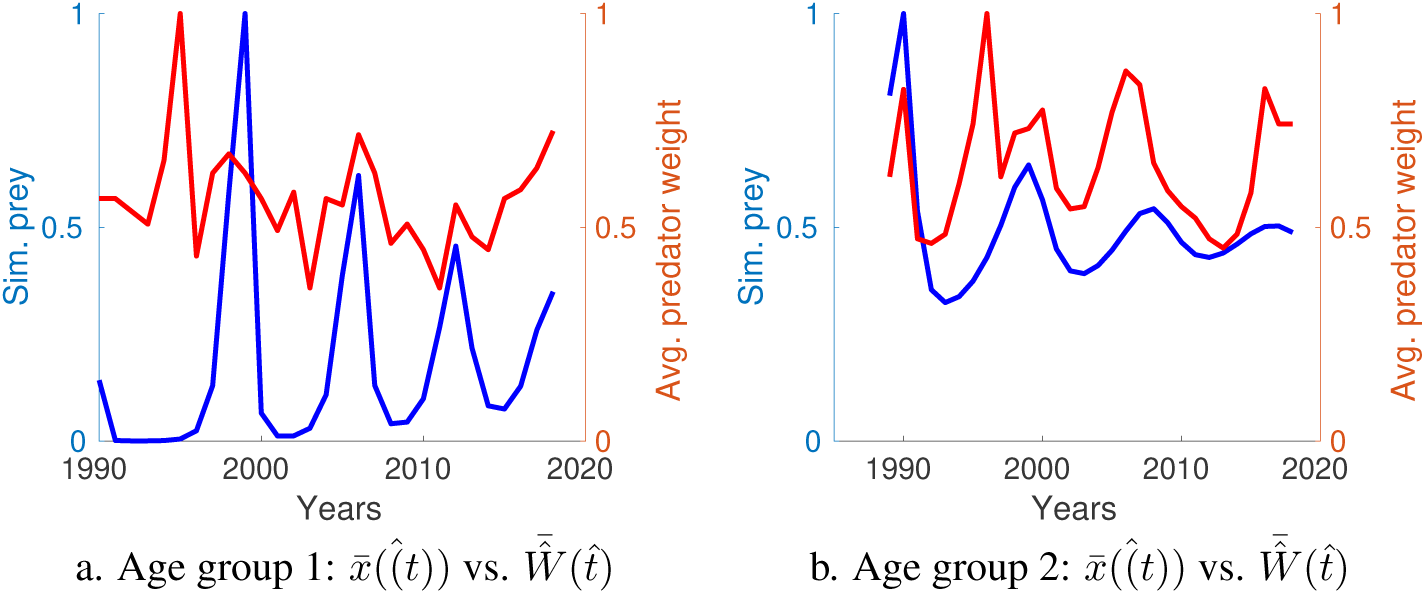
Scaled total biomass of prey, 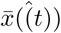 and scaled predator weight, 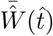 **Left:** 1-year old prey in the period 1990–2018 (a.), **Right:** 2-year old prey in the period 1989–2018 (b.)

